# Low-grade systemic inflammation stimulates microglial turnover and accelerates the onset of Alzheimer’s-like pathology

**DOI:** 10.1101/2024.03.15.585224

**Authors:** Monica Guerrero-Carrasco, Imogen Targett, Adrian Olmos-Alonso, Mariana Vargas-Caballero, Diego Gomez-Nicola

**Affiliations:** School of Biological Sciences, University of Southampton, Southampton General Hospital, United Kingdom; Institute for Life Sciences (IfLS), University of Southampton, United Kingdom

## Abstract

Several *in vivo* studies have shown that systemic inflammation, mimicked by LPS, triggers an inflammatory response in the CNS, driven by microglia, characterised by an increase in inflammatory cytokines and associated sickness behaviour. However, most studies induce relatively high systemic inflammation, not directly compared with the more common low grade inflammatory events experienced in humans during the life course. Using mice, we investigated the effects of low-grade systemic inflammation during an otherwise healthy early life, and how this may pre-condition the onset and severity of Alzheimer’s disease (AD)-like pathology. Our results indicate that low grade systemic inflammation induces sub-threshold brain inflammation and promotes microglial proliferation driven by the CSF1R pathway, contrary to the effects caused by high systemic inflammation. In addition, repeated systemic challenges with low grade LPS induce disease-associated microglia. Finally, using an inducible model of AD-like pathology (Line 102 mice), we observed that pre-conditioning with repeated doses of low-grade systemic inflammation, prior to APP induction, promotes a detrimental effect later in life, leading to an increase in Aβ accumulation and disease-associated microglia. These results support the notion that episodic low grade systemic inflammation has the potential to influence the onset and severity of age-related neurological disorders, such as Alzheimer’s disease.

## INTRODUCTION

Microglial cells are the main tissue-resident immune cells of the brain. They are specialised myeloid cells, derived from erythro-myeloid progenitors originated in the yolk sac during early embryogenesis (Ginhoux et al., 2010; Mass et al., 2016). The microglial population is maintained by self-renewal turning over multiple times during a mouse and human lifetime (Askew et al., 2017; Menassa et al., 2022). Microglial turnover is regulated by signalling through CSF1R (Askew et al., 2017), a tyrosine kinase receptor encoded by the c-fms proto-oncogene (Yeung and Stanley, 2003). This receptor is activated by two alternative ligands, CSF1 and IL-34 (Lin et al., 2008; Pixley and Stanley, 2004; Sherr et al., 1985), with key roles in microglial viability and proliferation (Greter et al., 2012; Kondo et al., 2007; Wegiel et al., 1998). Pharmacological inhibition of the activity of CSF1R leads to a reduction of microglial numbers in different animal models of neurodegenerative disease (Gomez-Nicola et al., 2013; Olmos-Alonso et al., 2016), associated with an amelioration of the AD-like pathology. Recently, we identified that the accumulated cycling events are linked to the onset of replicative senescence in microglia, a mechanism that accelerates early AD-like pathology in mice (Hu et al., 2021).

Systemic inflammation has been widely demonstrated to have a large impact in the inflammatory response within the CNS, in mice and humans (Holmes et al., 2009; Perry et al., 2007; Püntener et al., 2012). In rodents, systemic inflammation triggered by LPS increases inflammatory cytokines in the brain, activates microglia and triggers sickness behaviour (Perry et al., 2007; Pfeiffer, 1892; Püntener et al., 2012; Riester et al., 2020). In addition, LPS can trigger a proliferative response in microglia in a dose dependent manner (Fukushima et al., 2015; Furube et al., 2018; Shankaran et al., 2007). These studies show that LPS has an effect in microglial proliferation, however it is still not clear if repeated doses have a similar effect than one dose, and if this increased proliferative response might be triggering a different phenotype in microglial cells. Wendeln et al. 2018 studied the effect of repeated systemic doses of LPS, observing immune tolerance after the second systemic dose. Upon the third and fourth dose the cytokine levels in the blood and brain were significantly reduced except for IL-10 supporting immune tolerance (Wendeln et al., 2018). In this study, and in others, the doses of LPS were relatively high, comparable to humans displaying an exaggerated inflammatory response, such a flu, daily. However, here we are interested in studying a paradigm mimicking milder inflammation, akin having cold episodes spaced for longer periods of time.

Several studies in mouse models of neurodegeneration have strongly suggested that systemic inflammation exacerbates the progression of neurodegenerative diseases by increasing the production of pro-inflammatory cytokines, promoting cell death, and exacerbating chronic activation of microglia (Furube et al., 2018). During neurodegeneration mice challenged with systemic inflammation display an exacerbated neuroinflammatory response, indicating that microglia are primed, trained, by ongoing neurodegeneration (Cunningham et al., 2005; Kitazawa et al., 2005; Murray et al., 2012; Qin et al., 2007; Wendeln et al., 2018). For humans, this mechanistic model is directly relevant to systemic inflammatory events developed after neurodegeneration has started, but it may not apply to the effect of repeated, low grade, inflammatory events that humans experience during a healthy life course, prior to the onset of neurodegeneration. Several lines of evidence indicate that neuroinflammation is a causal factor for AD, contributing as much as the plaques and neurofibrillary tangles to the pathogenesis (Heneka et al., 2015). GWAS studies have identified genes related to innate immunity as risk genes of late onset of AD (Webers et al., 2020). In mice, there is some evidence suggesting that systemic insults might be triggering AD-like pathology later in life. Administration of Poly I:C, a viral mimetic, to pregnant dams, followed by one systemic injection in adulthood, was shown in wild type mice to induce signs of Aβ accumulation (Krstic et al., 2012).

Hence, we now hypothesise that low-grade inflammatory challenges, prior to the onset of Alzheimer’s disease, would drive an accelerated turnover of microglia, pre-conditioning the cells to develop an aggravated response to the onset of Aβ accumulation in the brain. Our results support that low grade systemic inflammation induces a transient wave of microglial proliferation, and that this is not observed when challenging in the presence of high-grade systemic inflammation. When low-grade, repeated, systemic inflammation precedes the onset of amyloid pathology, the disease progressed faster, with elevated indicators of AD-like pathology. In sum, our results support the hypothesis that repeated low-grade systemic inflammation preconditions microglia to subsequent AD-like pathology, accelerating the onset of the pathology.

## METHODS

### Experimental animals

C57BL/6, c-fms EGFP (Macgreen) (Sasmono et al., 2003) and TetO-APPSweInd (referred to as APP/tTA, Line 102) mice (Jankowsky et al., 2005) were bred and maintained at the Biomedical Research Facility (BRF; University of Southampton, UK). Both Macgreen and Line 102 mice were bred in a C57BL/6 background. Colonies of MacGreen mice were maintained as homozygotes. We used TetO-APPSweInd mice to model AD. This model expresses a chimeric mouse/human APP695 with the Swedish (KM570/571NL) and Indiana (V617F) mutation. The APP695swe/Ind sequence is downstream of the tetracycline (tetO) promoter and mouse prion protein exons 1-2. Hence, APP expression is regulated by a tetracycline-controlled transactivator protein (tTa) under the control of a tissue-specific promoter. The expression of tTA is inhibited while mice were fed with doxycycline (DOX), preventing the expression of APP. However, when DOX is withdrawn from the diet, mice express tTA and, consequently, APP, starting to accumulate Aβ in the cortex and hippocampus (Jankowsky et al., 2005; Sri et al., 2019). To maintain the colony male or female APP+/tTA-were crossed with male or female APP-/tTA+ to generate double transgenic APP/tTA mice.

C57BL/6, and Macgreen mice were provided with a standard RM1 diet (RM1, SDS UK) and water *ad libitum*. APP/tTA mice were provided with doxycycline diet (DOX; 625mg/Kg; TestDiet Limited) (Sri et al., 2019), and water *ad libitum.* All experimental procedures were approved by a local ethical review committee and conducted in accordance with relevant personal and project licenses under the UK Animals (Scientific Procedures) Act (1986). All mice were housed in groups of two or three, unless stated otherwise, per cage with wood chip bedding and maintained under a standard light-dark cycle (12:12-hour light-dark cycle with the light on at 7 am), at a temperature of 19-23°C and a humidity of 45-65% as per Home Office requirements.

### Systemic inflammatory challenge with LPS

LPS from Salmonella Enterica (serotype abortus equi; Sigma Aldrich, UK) (1mg/kg or 0.1mg/kg, 0.1ml/10g weight in sterile saline) or saline was injected intraperitoneally in mice (all groups were a mix of male and female mice).

For the experiment of single systemic challenge with LPS 2-3 months old C57BL/6 or Macgreen mice were divided into three groups (control, LPS-low and LPS-high). Control mice were injected with saline, LPS-low mice were injected with 0.1mg/kg LPS, and LPS-high mice were injected with 1mg/kg LPS. After the injection, mice were kept in individual cages, in order to measure individual burrowing behaviour. In addition, to analyse microglial proliferation, mice were injected with BrdU (Sigma-Aldrich; IP; 75mg/Kg, 0.1ml/10g weight in sterile saline) 18 hours before culling. Finally, mice were culled 48h or 72h after the LPS or saline injection.

### Inhibition of microglia proliferation

In order to inhibit microglial proliferation, 3-month-old C57BL/6 mice were treated with the CSF1R inhibitor GW2580 by oral gavage. GW2580 was diluted in 0.5% hydroxypropylmethylcellulose and 0.1% Tween 80 and dosed by oral gavage daily at 0.2mL per mouse (75mg/kg) throughout the experiment (Olmos-Alonso et al., 2016). Mice were divided in 4 groups: Control, Control GW2580, LPS-low, LPS-low+GW2580, all groups were a mix of female and male mice. The control and control GW2580 groups were both injected i.p with saline on day 3, additionally, the control GW2580 was treated with GW2580 by oral gavage daily. The LPS-low and LPS-low GW2580 groups were both i.p injected with 0.1mg/Kg LPS on day 3, with the LPS-low GW2580 treated by oral gavage daily throughout the experiment. Finally, to be able to analyse microglial proliferation all groups were i.p injected with BrdU (Sigma-Aldrich; IP; 75mg/Kg, 0.1ml/10g weight in sterile saline) 18 hours before culling on day 6.

### Repeated challenges with systemic LPS

3-month-old C57BL/6 mice were divided into 6 experimental groups, all groups were a mix of female and male mice. There were three control groups injected i.p with 1, 2 or 3 doses of saline spaced one week apart and three LPS-low groups, injected i.p with 1, 2 or 3 doses of 0.1mg/kg LPS, spaced one week apart.

### LPS and AD-like model

AD was modelled by using APP/tTA mice. First, we did a pilot experiment to study if DOX, alone, would affect the effect of LPS on microglial cells. Mice were injected with 0.1mg/Kg LPS or saline and were provided with either DOX or control diet. Sickness behaviour was measured on the day of the injection and the following day. All groups were a mix of female and male mice, except Saline + DOX that were only female mice Mice were fed with DOX from conception to weaning and throughout the experiment, receiving 3 i.p injections of saline or 0.1mg/kg LPS spaced one week apart. One week after the last i.p, on week 4, the DOX diet was withdrawn and replaced with regular RM1 diet for 16 weeks to allow tTA expression and switch-on the APP transgene, resulting in accumulation of Aβ-plaques. After 16 weeks of regular diet, mice were culled and samples were collected. All groups were a mix of female and male mice, except the group APP/tTA treated with saline that were all male.

### Sickness behaviour

Plastic cylinders (20cm long 6.8cm diameter) were filled with standard diet food pellets (RM1) and weighed and placed in individual mouse cages for two hours or overnight. After 2h or 18h, the remaining pellets within the cylinder were weighed to calculate the amount of displaced food (“burrowed”). Then mice were returned to their home cages or, if they stayed overnight, then they remained in the same cage. We also monitored weight loss to assess sickness behaviour the first three days after the i.p injection.

### Tissue extraction and processing

Mice were terminally anaesthetised with an overdose of pentobarbitone sodium i.p injected (20% weight/volume; AnimalCare, UK) and then perfused with 20mL of 0.9% saline and heparin (1/1000; Wockhardt, UK). Following perfusion, the brain was extracted and divided into two hemispheres. One of the hemispheres was snap frozen (−80°C) to be later used to measure cytokine concentration and RNA expression. The other hemisphere was processed for histology, postfixing in 4% PFA for 3 hours and then in 2% PFA overnight at 4°C. The following day, the hemi-brains were transferred to PBS and kept at 4°C until processing for sectioning using a vibrating microtome at 35μm thickness. Alternatively, they were included cryosprotected with sucrose, and included in OCT and stored at –20°C until they were sectioned on the cryostat at 35μm thickness. Later, the sections were kept at −20°C in cryoprotectant (300 g Sucrose + 10 g Polyvinyl-pyrrolidone (PVP-40) + 300 mL Ethylene glycol + 500 mL 0.1M phosphate buffer) until they were analysed in free floating.

Also, blood was collected by cardiac puncture to assess the immune response in the periphery by measuring the concentration of circulating cytokines. After collection of the blood, by an incision in the heart’s atrium, into BD Microtainer® K2E tubes (0.1mg K2-EDTA additive; Becton Dickinson, USA), blood was left at room temperature to allow coagulation. Later, it was centrifuged at 12000g for 10 minutes at 4°C and the serum was collected in a different Eppendorf and stored at −80°C.

### Immunohistochemistry

For brightfield microscopy, the endogenous peroxidase and phosphatase activity were blocked with Dual Endogenous Enzyme Block (Dako, Agilent Technologies, USA) for 15 minutes and then washed with PBST0.1% (3 x 5 minutes; PBS + 0.1% Tween20). Then, the tissue was incubated with blocking solution (PBS 0.2% TritonX100 or Tween20 + 5% BSA + 5% serum of host animal of the secondary antibody) to prevent unspecific binding of the antibodies. Sections were incubated with the following primary antibodies, diluted in blocking solution, overnight at 4°C: Chicken anti-GFP (1/500, Aves lab 1010); Rat anti-BrdU (1/500, Abcam ab2626); Rabbit Anti-Iba (1/500, In-House Covalab, Imai et al., 1996); Rat Anti-Dectin-1 (1/200, InvivoGen mabg-mdect); Mouse Anti-6E10 (1/500, Biolegend, sig39320). Sections were then washed and incubated with secondary biotinylated antibodies, diluted in Impress Kit (Vector Laboratories) or Alexa488/Alexa 568 conjugated secondary antibodies (Invitrogen). For immunofluorescence, the tissue was washed with PB (0.1M phosphate buffer; 3 x 5 minutes) and mounted on gelatine coated slides and coverslipped with Mowiol/DABCO. For brightfield immunohistochemistry, sections were washed and incubated with avidin-biotin complex (ABC) kit and signal was developed with peroxidase substrate diaminobenzidine (DAB) and/or BCIP/NBT kit (Vector) following manufacturer instructions. For immunofluorescence, sections were mounted on gelatinized or superfrost slides and coverslipped with Mowiol/DABCO. For brightfield immunohistochemistry sections were dehydrated in IMS (70%, 90% and 100%; 2 minutes in each) to coverslipped with DPX (Sigma).

The general protocol was modified to detect BrdU. The sections were pre-treated with citrate buffer (2.1g/L, pH6) for antigen retrieval by boiling for 30 seconds, followed by 5 minutes in ice-cold citrate buffer, repeating the process 3 times. The sections were then washed and incubated for 30 minutes at 37°C with 2N HCl to expose the BrdU epitope.

The general protocol was modified to detect human Aβ with the 6E10 antibody. The sections were pre-treated with formic acid 70% for 15 minutes and after washing the sections were directly mounted/coverslipped before imaging.

### Staining Aβ plaques with Congo Red

Sections were stained with Congo Red after completion of immunohistochemical staining, prior to dehydrating and coverslipping. Sections were incubated with 1% Congo Red for 30 minutes at RT. Then, they were rinsed with dH2O and later briefly immersed in alkaline alcohol (1%NaOH in 50% EtOH) to differentiate. The sections were then rinsed in tap water for 3-5 minutes and finally dehydrated in IMS (70%, 90% and 100%; 2 minutes in each) and immersed in xylene (15 minutes) to be coverslipped with DPX as mounting medium.

### Image acquisition and analysis

Images of the hippocampus (CA1) and cortex were taken with a microscope. Fluorescence images were obtained with a Leica CTR DM5000B coupled to a Leica DFC300FX camera (LASAF software) or a laser confocal microscope Leica TSC-SP8 confocal system on inverted microscope frames (DMi 8) with the LAS-X acquisition software. Brightfield images were taken with a Leica DM4B coupled to a Leica DFC7000 T camera (LASF software).

Images (n=3 sections/mouse), were analysed with ImageJ (Schindelin et al. 2009). The antigen-positive cells were counted with the “Cell Counter” plugin. If analysing brightfield double immunostaining, the deconvolution tool was used to better distinguish double positives. For analysis, the cell density was calculated (cells/mm^2^), also calculating density of the double positives and the % of double positives. The proliferative index was also calculated, dividing the total number of Iba1^+^BrdU^+^ cells by the total number of Iba1^+^ cells (Iba1^+^+BrdU^+^ /Iba1^+^) x100). Additionally, the dectin-1^+^ staining was counted directly on the microscope (3 sections/mouse) from the whole CA1 area of the hippocampus in WT mice. In APP/tTA mouse the dectin-1^+^ cells were counted from the whole cortex area and around the plaques.

For analysing microglial morphology, images of microglial cells were taken in the TSC-SP8 confocal microscope. Microglial individual territory area was measured and analysed with ImageJ. The brightness and contrast of the images were adjusted, to detect as many branches of cells as possible. The area occupied by 10 microglial cells in the CA1 region and their soma was manually measured on three sections per mouse. This involved outlining each cell and its visible branches. Subsequently, the average area of the 30 microglial cells measured in each mouse was calculated to determine the average microglial cell territory per mouse. (Fernández-Arjona et al., 2017; Torres-Platas, G, S Comeau et al., 2014; Verdonk et al., 2016).

### Analysis of gene expression by reverse transcriptase qPCR

RNA from the rostral half of the hippocampus of C57BL/6 mice was extracted with TRIzol^®^ (Life Tecnologies). RNA was quantified with a Nanodrop (Thermo Scientific) and the iScript^TM^ cDNA Synthesis Kit (Bio-Rad) was used to reverse transcribe the RNA, following the manufacturer’s instructions. cDNA samples were analysed by RT-PCR with iTaq^TM^ Universal SYBR^®^ Green supermix (Bio-Rad) and the following gene-specific primers (Sigma-Aldrich): *csf1* (NM_007778.4; FW, agtattgccaaggaggtgtcag, RV, atctggcatgaagtctccattt), *csf1r* (NM_001037859.2; FW, gcagtaccaccatccacttgta, RV, gtgagacactgtccttcagtgc), *pu.1* (NM_011355.1; FW, cagaagggcaaccgcaagaa, RV, gccgctgaactggtaggtga), *cebpα* (NM_007678.3; FW, agcttacaacaggccaggtttc, RV, cggctggcgacatacagtac), *runx1* (NM_001111021; FW caggcaggacgaatcacact, RV, ctcgtgctggcatctctcat), *tnfα* (NM_013693; FW, aggcactcccccaaaagatg, RV, ttgctacgacgtgggctac), *il10* (NM_010548.2; FW ggcccagaaatcaaggagca, RV, acaggggagaaatcgatgacag), *il6* (NM_031168.1; FW, tagtccttcctaccccaatttcc, RV, ttggtccttagccactccttc), *il1β* (NM_008361.3; FW, gaaatgccaccttttgacagtg, RV, tggatgctctcatcaggacag), *il4* (NM_021283.2; FW, agcaacgaagaacaccacag, RV, gcatcgaaaagcccgaaagag), *gapdh* (NM_008084.2; FW, tgaacgggaagctcactgg, RV, tccaccaccctgttgctgta). Electrophoresis of the PCR product in a 1.5% agarose gel was used to check the quality of the primers and PCR reaction. The data was analysed using the 2^-ΔΔC(t)^ method (Livak and Schmittgen 2001), where the expression of the housekeeping gene (*gapdh*) is used as a reference and compared to the expression of the gene of interest.

### Protein extraction and quantification

The caudal half of the hippocampus was dissected, weighed, homogenised with a mechanical homogeniser and diluted 5 times with RIPA buffer (5ml of Pierce RIPA buffer, Thermo Scientific #89900), with protease inhibitor cocktail, Roche #04693124001) at 4°C. Protein concentration was quantified with a BCA protein assay kit (Pierce BCA, Thermo Scientific). Proteins were diluted in RIPA buffer to a concentration of 2.5mg/mL.

### Western Blot

The samples were diluted (3:1) with 4x sample buffer (8% SDS, 20% 2-mercaptoethanol, 40% glycerol, 0.008% bromophenol blue, in 0.250M Tris-HCl, pH 6.8) to load 30µg per sample. The samples were loaded and run in 7% SDS-polyacrylamide gels and separated through electrophoresis. Then, they were blotted onto a nitrocellulose membrane, followed by blocking with 1X Tris-Buffered Saline, 0.1% Tween 20 Detergent (TBS-T) + 5% BSA for 1h at room temperature and washed with TBS-T + 0.1%Tween-20 (3×5minutes). Then, the membrane was incubated overnight in primary antibody (mouse anti-6E10-Biolegend sig39320, 1/1000; rabbit anti-β-tubulin-Cell signalling 2128S, 1/1000) at 4°C, followed by washing and incubation with the appropriate secondary antibody (LICOR, 1/10000) for 1h at room temperature. The membrane was then washed and scanned with an Odyssey CLx Infrared Imaging System (LICOR) at the appropriate wavelength. Immunolabelled bands were quantified with the EmpiriaStudio software.

### Multiplex analysis of inflammatory cytokines

Following protein quantification, brain samples were used undiluted, with blood samples diluted 1/10. IFN-γ, IL-1β, IL-2, IL-4, IL-5, IL-6, IL12p70, IL-10, KC/GRO and TNF-α were analysed with the Proinflammatory Panel 1 (mouse) MSD kit (K15048-Series Mesoscale Discovery, USA) following the manufacturer’s instructions. Standards and samples were added to the 96-well plate, incubated and read using a QuickPlex SQ 120 plate reader (Mesoscale Discovery, USA). Worckbench 4.0 (Mesoscale Discovery, USA) was used to acquire and analyse data.

### Statistical analysis

Data from independent biological replicates were shown as mean±SEM, and analysed with GraphPad Prism 7 software package. For all datasets where normality and homoscedasticity assumptions were reached, we used one-way or two-way ANOVA followed by the Tukey *post hoc* test for multiple comparisons or a Sidák’s multiple comparisons test. When normality and homoscedasticity assumptions were not reached, the dataset was analysed using the Kruskal-Wallis test or Mann-Whitney test. Details of specific statistical test and N are displayed in the figure legends. Significant differences were considered when p<0.05.

## RESULTS

### LPS triggers a dose-dependent systemic and central inflammatory response

First, we wanted to compare the dose-dependency of the LPS-induced systemic and brain inflammatory response **(Figure 1A)**. Hence, we compared two different doses of LPS: 0.1mg/Kg (LPS-low), mimicking low grade systemic inflammation (Skelly et al., 2013) and 1mg/kg (LPS-high), which triggers an acute inflammatory response (Furube et al., 2018). 24h after the systemic injection, there is a significant decrease in burrowing behaviour in animals injected with LPS, independently of the dose **(Figure 1B)**, although the LPS-low group displayed a milder phenotype. The sickness behaviour induced by LPS is accompanied by weight loss, which increases over time in the LPS-high group, while the LPS-low group present a very mild change in weight **(Figure 1C)**.

**Figure 1.**
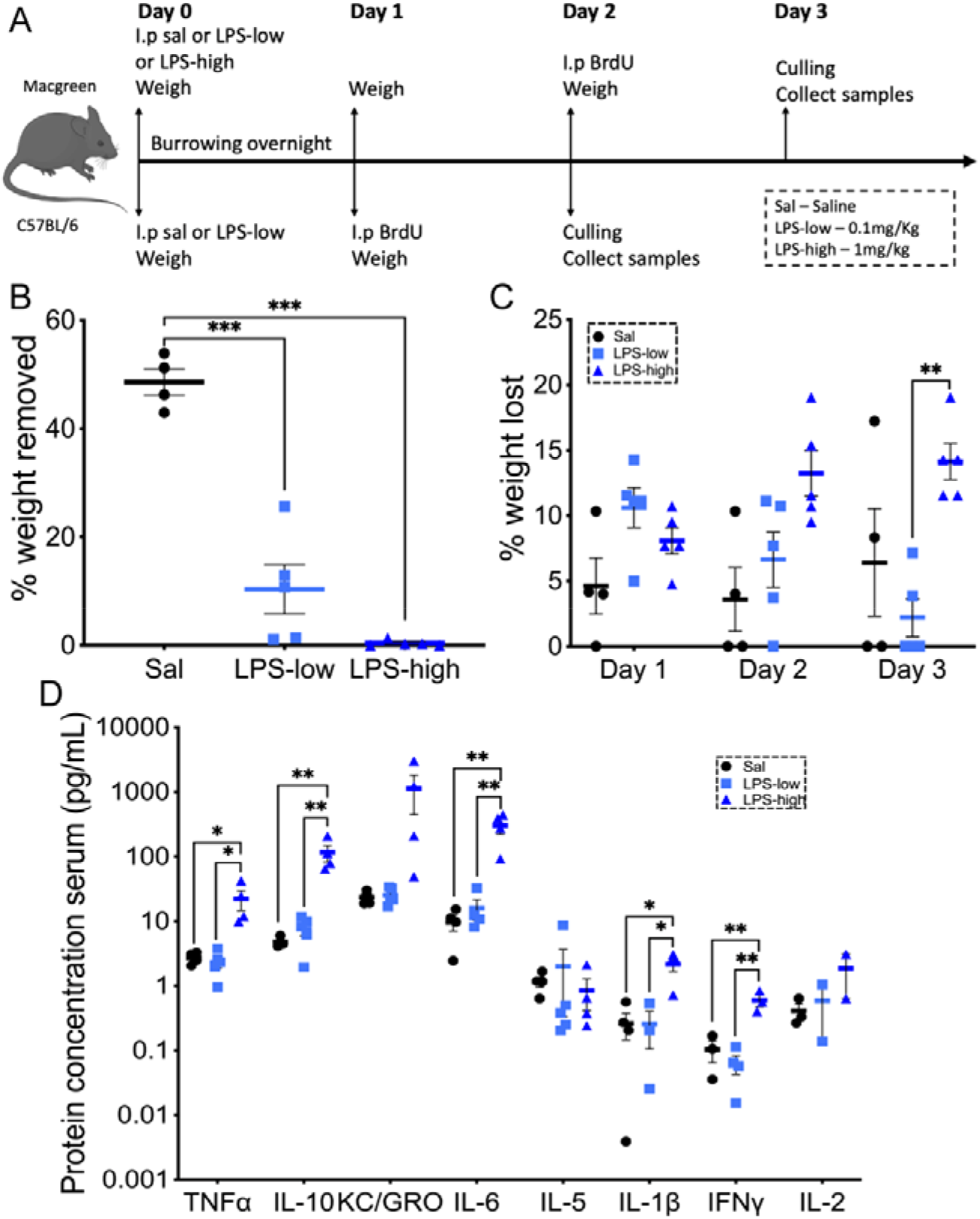
LPS-high triggers a sustained sickness behaviour while LPS-low triggers a mild sickness behaviour in 3 months old mice. (A) Experimental plan to study LPS as a model of systemic inflammation (B) Burrowing behaviour measured overnight from day 0 to day 1 after i.p injection of Sal (n=4), (0.1mg/kg LPS (LPS-low; n=5) or 1mg/Kg LPS (LPS-high; n=5). Data were analysed with a One-way ANOVA and a post hoc Tukey’s test. (C) Percentage of weight lost in each of the three days of the experiment respectively to the baseline weight at day 0. Data were analysed with a 2-way ANOVA and a post hoc Tukey’s test. Data shown represented as mean±SEM. (D) Mesoscale analysis of TNFα, KC/GRO, IL-6, IL-5, IL-1β, IFNγ, IL-2 and IL-10 in the serum of MacGreen mice after i.p injection of Saline (Sal; n=4) or 0.1mg/Kg LPS (LPS-low; n=4-5) or 1mg/kg LPS (LPS-high; n=4-5) at 3 days post-injection. TNFα, IL-10, IL-6, IL-1β, IFNγ and IL-2 analysed with One-Way ANOVA and a post-hoc Tukey’s test, IL-5 analysed with Kruskal-Wallis test and Dunn’s multiple comparison test. Data shown represented as mean±SEM in log scale. Statistical differences *p<0.05, **p<0.01, ***p<0.001.

The systemic inflammatory response after challenge with LPS-high is significantly higher when compared with saline or LPS-low **(Figure 1D)**. We detected a significant increase in the concentration of TNFα, IL-1β, IL-6, IL-10 and INFγ in the serum of mice treated with systemic LPS-high compared with Sal and LPS-low **(Figure 1D)**.These results are in agreement with previous studies showing that systemic doses of LPS of 0.5mg/kg or higher trigger an acute inflammatory response (Biesmans et al., 2013; Erickson and Banks, 2011; Wendeln et al., 2018). However, we observed no significant differences after challenge with LPS-low, when compared to control mice **(Figure 1D).**

These results support that LPS-low triggers mild and transient systemic inflammation and sickness behaviour, with faster recovery when compared to LPS-high, supporting the idea that LPS-low is an optimal model of low-grade systemic inflammation.

We then monitored the effect of systemic LPS challenge on the inflammatory response in the hippocampus. At day 3 after systemic LPS challenge, we found an elevated mRNA expression of *Il-6*, *Il-10*, *Il-1β* and *Tnfα* in the LPS-high group, with no changes observed after LPS-low **(Figure 2A)**. At the protein level, both IL-1β and TNFα were found elevated after LPS-high, with no changes observed after LPS-low **(Figure 2B)**. These results indicate that LPS-high triggers a prominent central inflammatory response that lasts at least 3 days, while LPS-low does not. The protein concentrations of IL-6, IFNγ, IL-2, IL-12p70, IL-5 or IL-10, IL-4 were found unchanged in all the different experimental groups, suggesting part of this response is already resolved by day 3.

**Figure 2.**
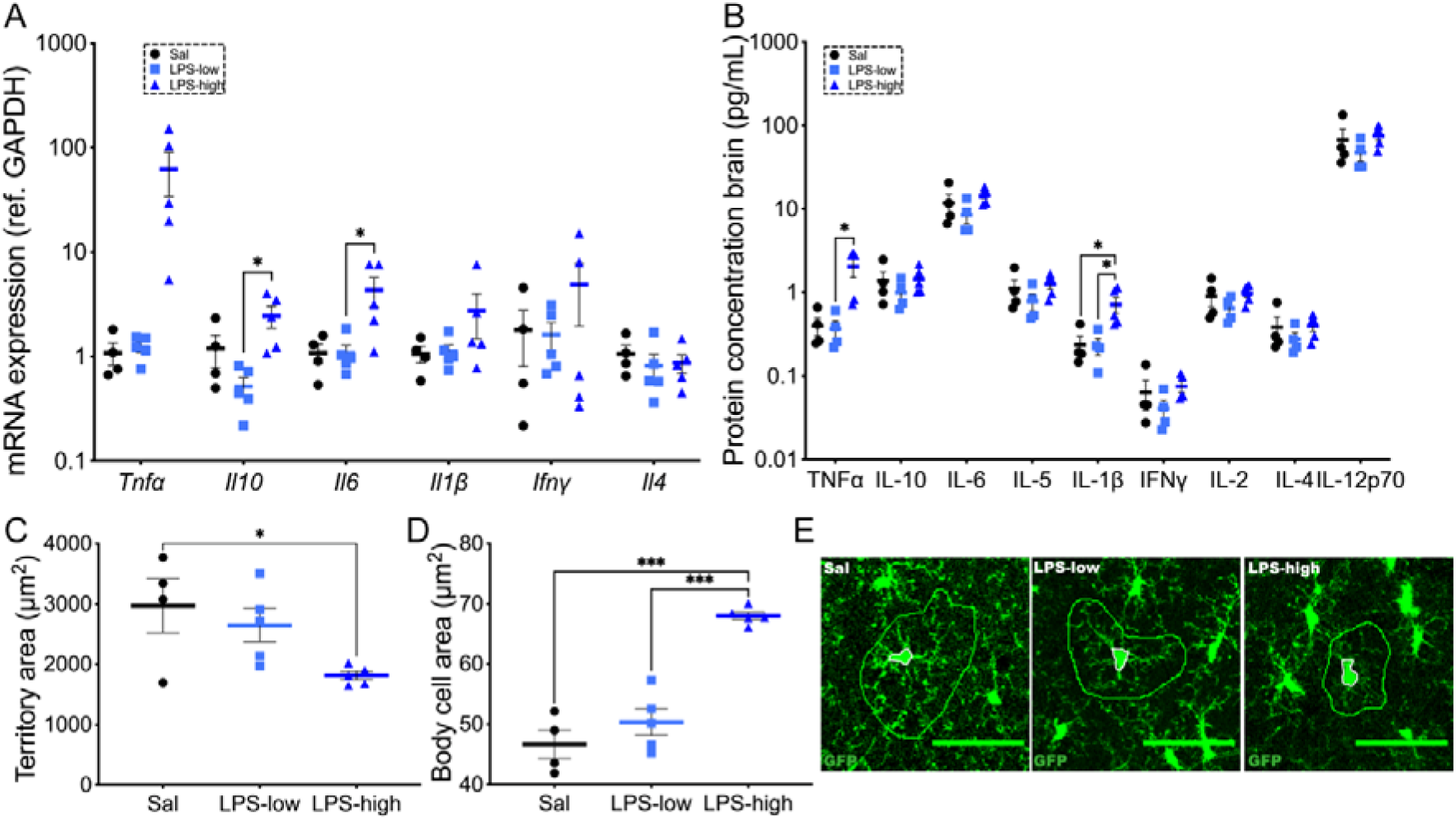
Inflammatory response in the hippocampus after challenge with LPS-high is significantly increased compared to Sal and LPS-low treatment. (A) RT-PCR analysis of the mRNA expression of Tnfα, Il6, Il1β, Ifnγ, Il10 and Il4 in the hippocampus of MacGreen mice after i.p injection of Saline (Sal; n=4) or 0.1mg/Kg LPS (LPS-low; n=5) or 1mg/kg LPS (LPS-high; n=5) at 3 days post-injection. Data analysed with One-way ANOVA and a post-hoc Tukey’s test (B) Mesoscale analysis of TNFα, IL-6, IL-5, IL-1β, IFNγ, IL-2, IL-12p70, IL-10 and IL-4 in the hippocampus of MacGreen mice after i.p injection of saline (Sal; n=4) or 0.1mg/Kg LPS (LPS-low; n=4-5) or 1mg/kg LPS (LPS-high; n=4-5) at 3 days post-injection TNFα analysed with Kruskal-Wallis and a Dunn’s multiple comparison IL-10, IL-6, IL-5, IL-1β, IFNγ, IL-2, IL-4, IL-12p70 analysed with One-way ANOVA and a post-hoc Tukey’s test. Data shown represented as mean±SEM in log scale. (C) Quantification of microglial territory area in the hippocampus (CA1) of MacGreen mice (n=4-5). (D) Quantification of microglial soma area in the hippocampus (CA1) of MacGreen mice (n=4-5) (E) Representative images of GFP^+^ microglial territories measured and traced in 30μm z-stacks in the CA1 hippocampal area from MacGreen mice i.p injected with saline (Sal), 0.1mg/Kg (LPS-low) or 1mg/kg (LPS-high). Data were analysed with a One-way ANOVA and a post hoc Tukey’s test. GFP^+^ in green. Scale bar 50μm. Data shown represented as mean±SEM. Statistical differences *p<0.05 and ***p<0.001.

We measured the morphological changes in microglia in response to LPS-low and LPS-high, as an indirect measure of activation. Activated microglia usually have retracted and thicker branches with a bigger soma, occupying a smaller territory area (Fernández-Arjona et al., 2017; Karperien et al., 2008; Kondo et al., 2011; Verdonk et al., 2016). On the contrary, homeostatic microglia tend to display a smaller soma with longer branches (Kondo et al., 2011; Verdonk et al., 2016). Hence, we measured the territory area and the soma area of microglial cells in CA1, 3 days after systemic inflammation **(Figure 2E)**. We observed a significant decrease in the territory area of microglia after systemic challenge with LPS-high compared with saline, with no significant difference detected in the LPS-low group **(Figure 2C)**. We also observed a significant increase in the microglial soma area after systemic challenge with LPS-high compared with saline and with LPS-low.

Overall, these data support that LPS-low drives a subthreshold inflammatory response in the brain, with minor impact on microglial activation.

### Low grade systemic inflammation (LPS-low) triggers a microglial proliferative response

We aimed to further explore the effects of systemic LPS on microglial cells, specifically analysing their proliferation.

We studied microglial proliferation at day 3 after systemic challenge with saline, LPS-low or LPS-high, by administering BrdU 18h prior to sample collection **(Figure 3A)**. The results showed a significant increase in microglial proliferation in mice challenged with LPS-low compared with control mice, and this effect is not observed after challenge with LPS-high **(Figure 3C)**. This increased proliferation did not impact the total microglial density although we observed a slight decrease in mice treated with LPS-high **(Figure 3B)**.

**Figure 3.**
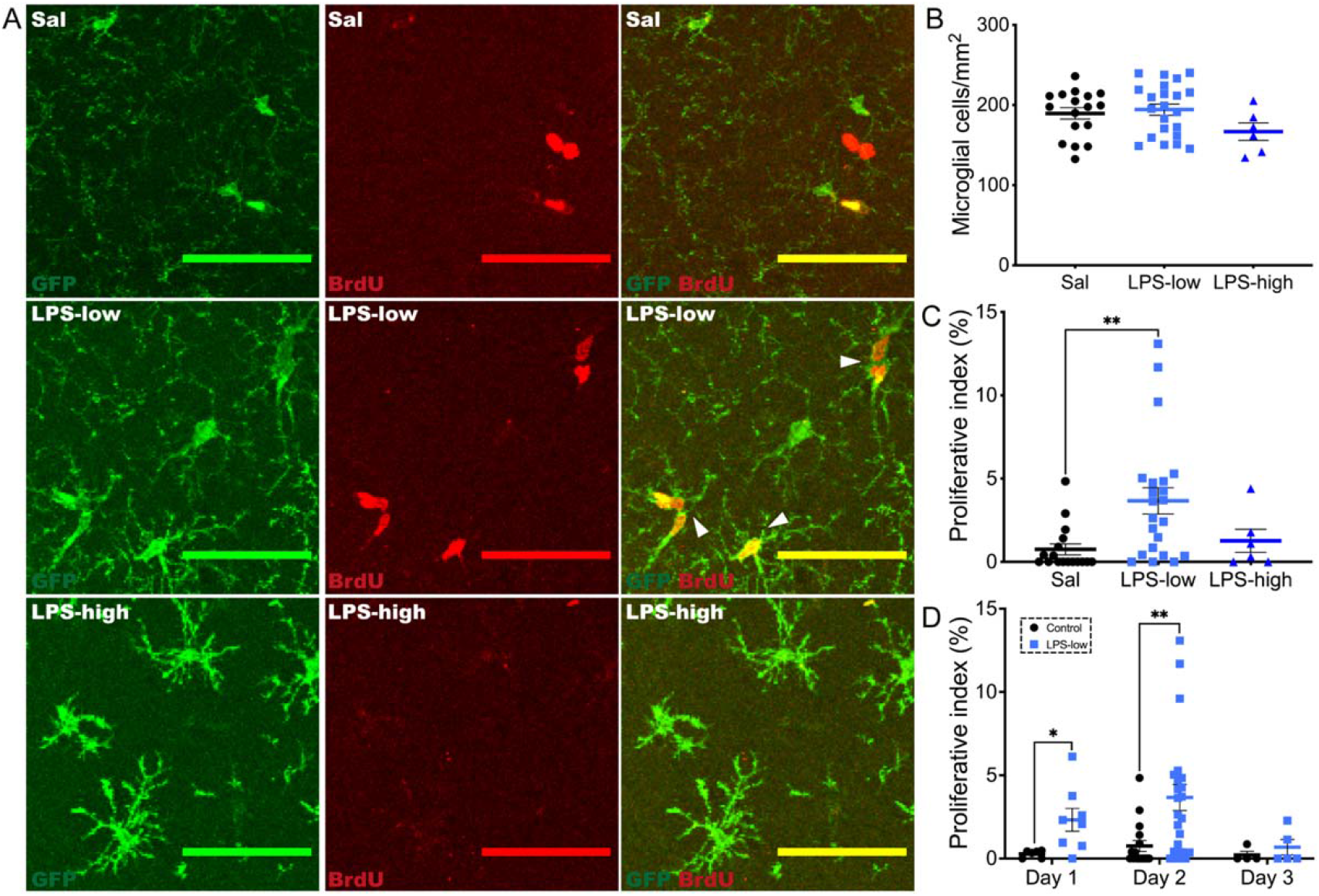
Microglial proliferation increases after systemic challenge with LPS-low. (A) Representative images of the hippocampus (CA1) of MacGreen mice systemically injected with Saline (Sal) or 0.1mg/Kg (LPS-low) or 1mg/kg (LPS-high). GFP^+^ or Iba1^+^ (Green) and BrdU^+^ (Red). White arrows indicate proliferating microglia. Scale bar 50μm. (B) Quantification of microglial density (GFP^+^ or Iba1^+^ cells/mm2) of mice injected with Sal (n=17) or LPS-low (n=22) or LPS-high (n=6) at 3 days post-injection. Data analysed with One-way ANOVA and a post-hoc Tukey’s test. (C) Quantification of the proliferative index (%) of Sal (n=17), LPS-low (n=22), and LPS-high (n=6) at 54h post-injection. Data analysed with Kruskal-Wallis with Dunn’s multiple comparisons test. (D) Quantification of the proliferative index (%) 30h post-injection of Sal (n=6) and LPS-low (n=8) and culled 2 days post-injection (day 1). 54h post-injection of Sal (n=17), LPS-low (n=22), culled 3 days post-injection (day 2). 72h post-injection of Sal (n=4), LPS-low (n=5), culled 3 days post-injection (day 3). Data analysed with unpaired t-test. Data shown represented as mean±SEM. Statistical differences *p<0.05, **p<0.01.

Then, we studied the time course of microglial proliferation after systemic injection of LPS-low, or saline, by administering BrdU 18h prior to sample collection on day 2 and on day 3 **(Figure 1A)**. We observed a significant increase in the proliferation of microglial cells in the CA1 of mice after LPS-low, peaking at day 2, compared with saline, and resolved by day 3 **(Figure 3D)**.

### Microglial proliferation after LPS-low is dependent on CSF1R activation

We studied the CSF1R-PU.1-CEBP/a axis, postulated to be the primary regulator of microglial proliferation in health and disease (Gomez-Nicola et al., 2013; Olmos-Alonso et al., 2016) **(Figure 4A)**. On day 2, we found a significant increase in the mRNA expression of *Csf1r and Pu.1* after LPS-low, compared with saline **(Figure 4B)**. The expression of other relevant genes, *Csf1*, *Cebpa* and *Runx1*, was found unchanged with the treatment **(Figure 4B)**. By day 3, the expression levels of the components of the CSF1R pathway returned to baseline **(Figure 4C)**, supporting our previous evidence that the increased microglial proliferation is resolved by day 3.

**Figure 4.**
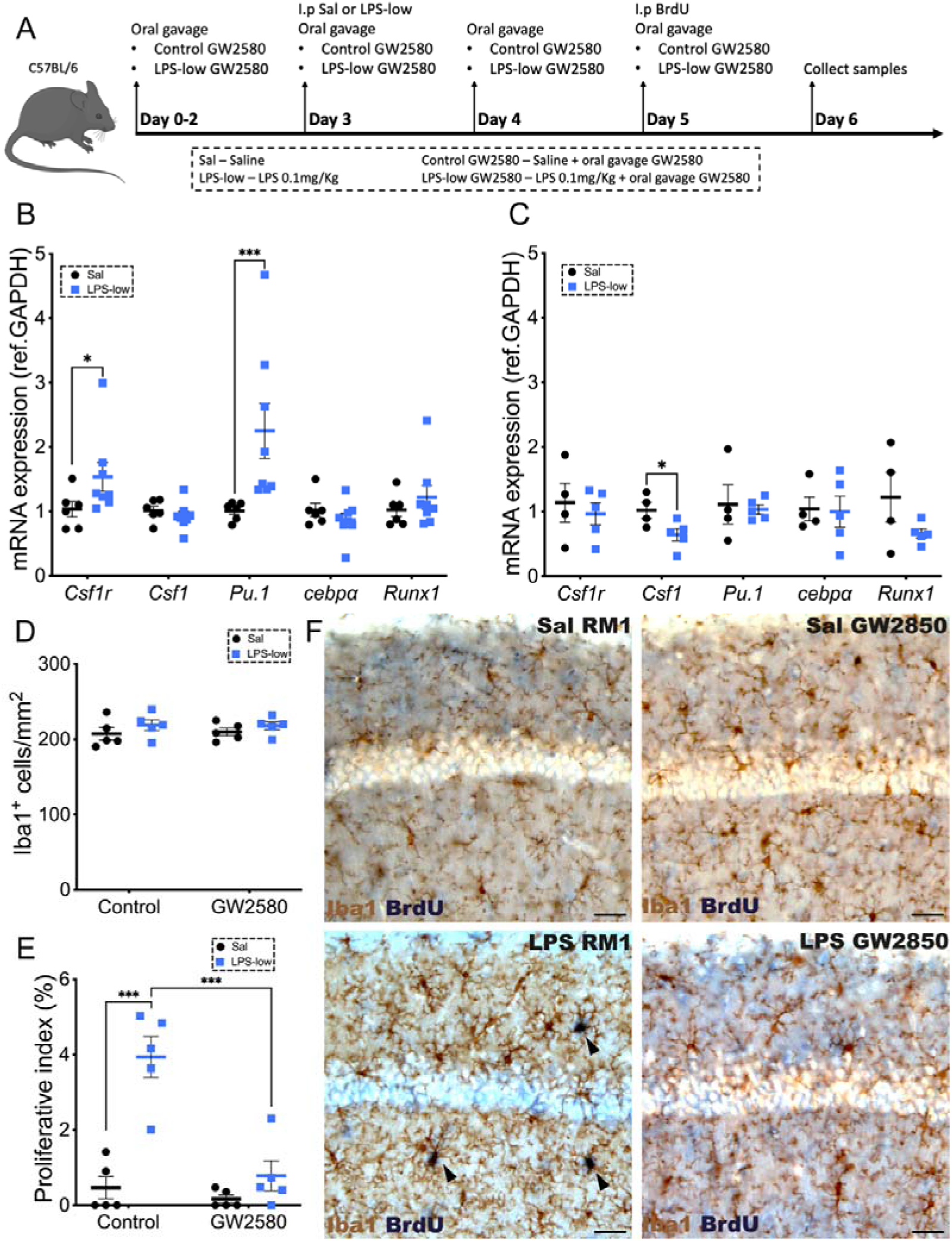
CSF1R pathway promotes microglial proliferation of LPS-low systemic injection. (A) Experimental plan to inhibit microglial proliferation (B, C) RT-PCR analysis of the mRNA expression of the components of the CSF1R-PU.1-CEBP/a axis (Csf1r, Csf1, Pu.1, cebpa and Runx1) after i.p injection of saline (Sal; n=6) or 0.1mg/Kg LPS (LPS-low; n=8) at 2 days post-injection (A: Csf1 analysed with unpaired t-test, Csf1r, Pu.1, cebpα, Runx1 analysed with Mann-Whitney test) or 3 days post-injection (B: data analysed with unpaired t-test). (D, E) Quantification of microglial density (Iba1^+^ cells/mm2 and proliferative index (%) in mice injected with Sal or LPS-low and fed with RM1 or GW2580 diet (n=5) at 3 days post-injection. Data analysed with a two-way ANOVA and a post hoc Tukey’s tests. (F) Representative images of the CA1 area of the hippocampus of mice i.p injected with Sal or LPS-low. Iba-1^+^ (brown in DAB) and BrdU^+^ (blue in AP) of mice fed with RM1 or GW2580. Black arrows indicate proliferating microglia. Scale bar 50μm. Statistical differences *p<0.05, and ***p<0.001. Data shown represented as mean±SEM.

Following the indication that the CSF1R pathway is activated after LPS-low, we tested its mechanistic role by using pharmacological inhibition of CSF1R with GW2580, an effective tyrosine kinase inhibitor (Askew et al., 2017; Olmos-Alonso et al., 2016). We dosed GW2580 by oral gavage daily (0.2mL per mouse, 75mg/Kg), starting 3 days prior to dosing saline or LPS-low, followed by administration of BrdU 18 hours before sample acquisition on 3 days after i.p injection of Sal or LPS **(Figure 4F)**. Microglial density was unaffected by treatment with GW2580 **(Figure 4D)**, consistent with our dosing regime and previous results (Askew et al., 2017; Hu et al., 2021; Olmos-Alonso et al., 2016). CSF1R blockade by GW2580 significantly decreased microglial proliferation in the hippocampus of mice treated with systemic LPS-low **(Figure 4E)**, confirming that LPS-low triggers microglial proliferation in a CSF1R-dependent manner.

### Repeated weekly doses of LPS-low trigger sickness behaviour and promote a change in the microglial phenotype

Repeated systemic doses of LPS can lead to immune tolerance, whereby microglial cells have an attenuated response to a second LPS stimulus (Liu et al., 2017; Wendeln et al., 2018). However, most reports that show immune tolerance to LPS use what we established as a high dose of LPS, and also consecutive daily challenges, overlapping with the inflammatory response from previous doses (Banasikowski et al., 2015; Liu et al., 2017; Wendeln et al., 2018). Hence, we wanted to study if repeated systemic challenges with LPS-low (3x LPS-low), spaced one week apart (to ensure resolution of inflammation), trigger immune tolerance in microglia. Analysis of sickness behaviour, studied by burrowing and weight, show that systemic 3x LPS-low seems not to trigger an overt immune tolerance, not observing tolerance by week 2 **(Figure 5A-B).** By the third week, the impact of systemic LPS-low on burrowing behaviour is attenuated, coinciding with a limited effect on weight loss, which might indicate a mild tolerance response **(Figure 5A-B)**.

**Figure 5.**
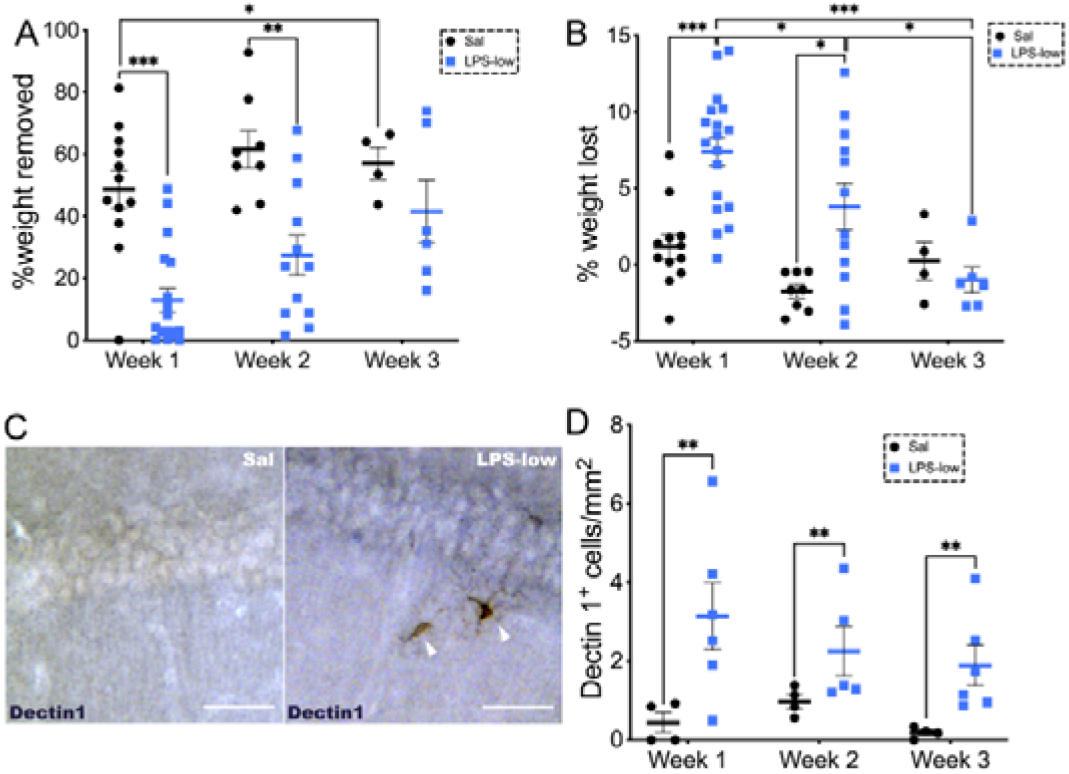
Repeated systemic dosage with LPS-low promotes mild sickness behaviour and dectin-1^+^. (A) Burrowing behaviour for 2 hours on day 0 after each weekly i.p injection of Sal (W1 n=12, W2 n=8, W3 n=4) or LPS-low (W1 n=18, W2 n=12; W3 n=6). (B) Percentage of weight lost 24h after i.p respectively to the base weight before i.p injection with Sal (W1 n=12, W2 n=8, W3 n=4) or LPS-low (W1 n=18, W2 n=12; W3 n=6). Data analysed with a 2-way ANOVA mixed-effect comparison Sidak’s multiple comparison test. (C) Representative images of CA1 of mice injected with 1 dose of 0.1mg/Kg LPS (LPS-low). White arrows show dectin1^+^ microglia. Dectin1^+^ in black (DAB + nickel ammonium). Scale bar 50μm. (D) Dectin1^+^ density in CA1 of mice systemically injection Sal 1, 2 or 3 doses (n=4) or 1, 2 or 3 LPS-low (n=5). Data analysed with 2-way ANOVA and a post-hoc Tukey’s test. Data shown represented as mean±SEM. Statistical differences *p<0.05, **p<0.01, ***p<0.001.

We followed to study if 3x LPS-low would alter the microglial phenotype, specifically inducing the dectin-1^+^ profile, which is a marker associated with an activation state present in AD-like pathology (Hu et al., 2021) **(Figure 5C)**. We observed a significant increase in the density of dectin-1^+^ cells in the CA1 of mice systemically injected with 1, 2 or 3 doses of LPS compared with saline **(Figure 5D)**. These results indicated that, in response to LPS-low, a fraction of the microglial population changed its phenotype to dectin-1^+^.

### Repeated exposure to LPS-low triggers sickness behaviour and affects A**β** accumulation in APP/tTA mice

In order to model the effect of repeated exposure to mild systemic inflammation in early life on the onset of AD-like pathology, we resourced to the APP/tTA mouse model (Han et al., 2012; Jankowsky et al., 2005). In presence of DOX, the APP transgene is silenced, allowing a *bona fide* pre-pathology window in which to model systemic inflammation. We first wanted to study if DOX, an antibiotic, would affect the effect of LPS-low on microglial cells, since previous studies had shown a decrease in hippocampal inflammation caused by LPS after treatment with DOX (Ferreira Mello et al., 2021, 2013). Thus, we compared mice systemically injected with Sal or LPS-low and fed with control diet or with DOX diet. DOX does not affect the sickness behaviour response observed in mice after dosing with LPS-low, evidenced by a significant difference in burrowing behaviour after LPS-low in both groups, fed with normal diet or fed with DOX diet **(Figure 6A)**. We also observed a significant difference in the weight loss between mice injected with Sal *vs* LPS-low in mice fed with normal diet or with DOX diet **(Figure 6B)**. These results support that a DOX-containing diet does not influence the responsiveness to systemic LPS challenge in APP/tTA mice.

**Figure 6.**
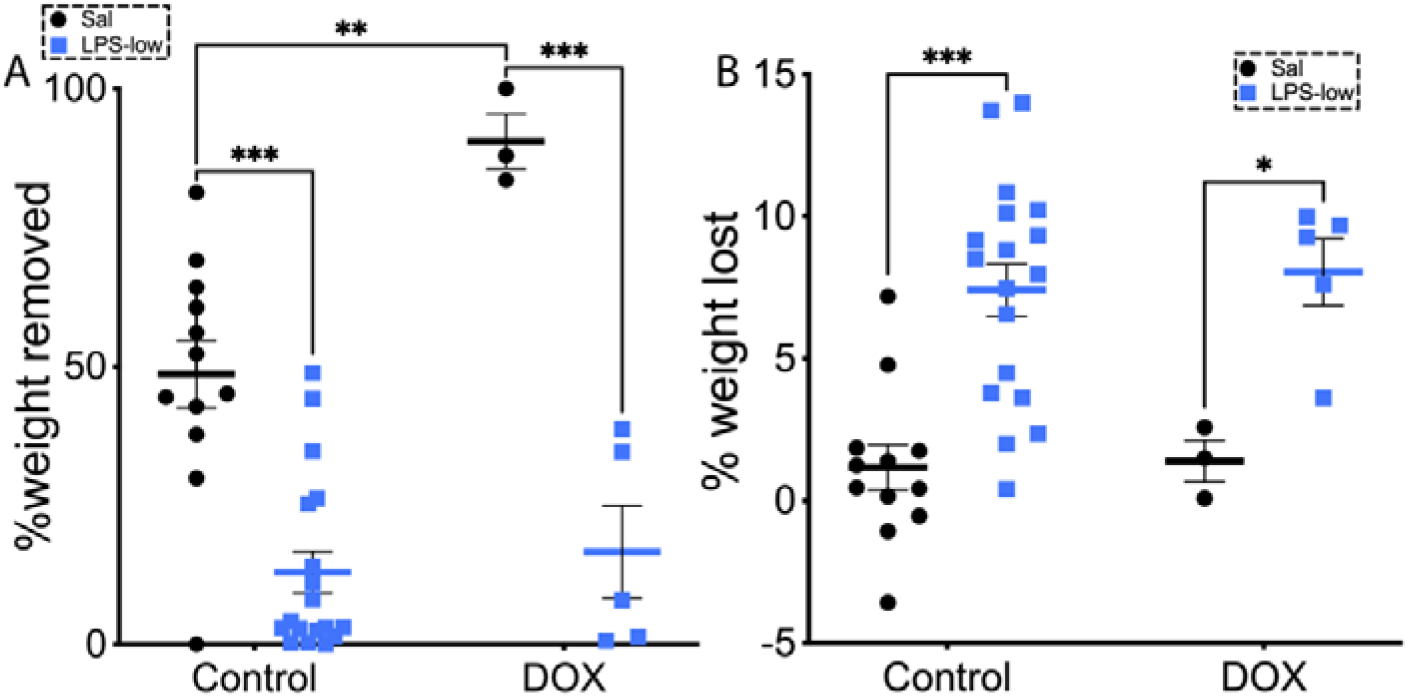
LPS-low triggers sickness behaviour in APP/tTA mice fed with DOX. (A) Burrowing behaviour for 2 hours on the day of the injection of mice fed with control and i.p injected with saline (Sal; n=12) or 0.1mg/Kg (LPS-low; n= 18) or mice fed with DOX and i.p injected with saline (Sal; n=3) or 0.1mg/Kg (LPS-low; n=5). (B) Percentage of weight lost 24h after i.p, related to the initial weight of mice fed with control and i.p injected with saline (Sal; n=12) or LPS-low (0.1mg/Kg LPS; n= 18) or mice fed with DOX and i.p injected with saline (Sal; n=3) or 0.1mg/Kg (LPS-low; n=5). Data analysed with 2-way ANOVA and Tukey’s multiple comparison test. Data shown represented as mean±SEM. Statistical differences ***p<0.001, **p<0.01, *p<0.05.

We next aimed to characterise the effect of our treatments on Aβ accumulation. In APP/tTA mice, there is a rapid increase of APP expression and Aβ levels following withdrawal of DOX (Sri et al., 2019). Hence, based on these studies, we kept mice in DOX from weaning and throughout the experiment, completing our 3x LPS-low during this transgene suppression period. We then withdrew DOX, and waited 16 weeks to analyse the density of Aβ plaques in the cortex, aiming to understand if 3x LPS-low had any impact on the onset of Aβ pathology. Our results show that pre-treatment with 3x LPS-low triggers a significantly higher accumulation of Aβ plaques in the cortex compared with mice treated with Sal **(Figure 7A,C)**. This increase was also observed in CA1, albeit not significant **(Figure 7B,E**). In addition, when measuring plaque burden in the cortex **(Figure 7D)** and CA1 area **(Figure 7F)**, there is an increasing trend in both areas in mice pre-treated with LPS-low compared with Sal (p=0.1).This effect is not due to modulation of the expression of the APP transgene, as detected by WB **(Figure 7G-H)**.

**Figure 7.**
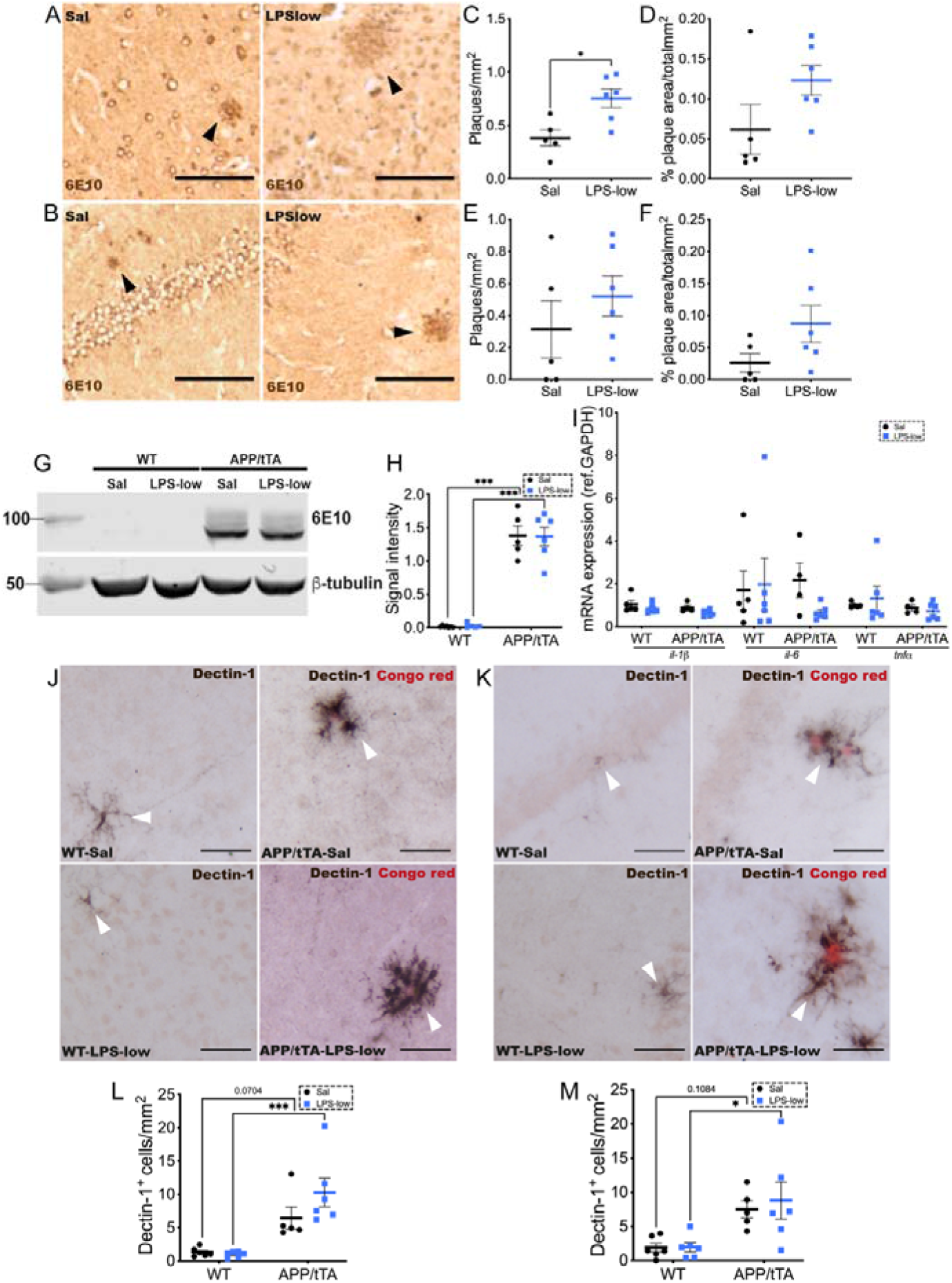
Pre-conditioning with repeated LPS-low promotes Aβ **accumulation and dectin-1^+^ in the cortex and CA1 of APP/tTa mice**. (A) Representative images of the cortex (B) CA1 area of APP/tTA mice 16 weeks off DOX after i.p injected 3 times with saline (Sal) or 0.1mg/Kg LPS (LPS-low). Scale bar 0.1mm. Black arrows indicate plaques (brown). (C) Plaque density (plaques/total area in mm2) in the cortex of APP/tTA mice i.p injected with Sal (n=5) or LPS-low (n=6). Data analysed with unpaired t-test. (D) Plaque load (%plaque area/ total area in mm^2^) in the cortex of APP/tTA mice i.p injected with Sal (n=5) or LPS-low (n=6). Data analysed with Mann-Whitney test. (E) Plaque density (plaques/total area in mm2) in the CA1 area of the hippocampus of APP/tTA mice i.p injected with Sal (n=5) or LPS-low (n=6). Data analysed with unpaired t-test. (F) Plaque burden (%plaque area/ total area in mm2) in the CA1 area of the hippocampus of APP/tTA mice i.p injected with Sal (n=5) or LPS-low (n=6). Data analysed with unpaired t-test. (G) Representative images of the WB bands of the 6E10 (APP) and its control (β-tubulin). (H) Signal intensity of the bands of 6E10 corrected by b-tubulin in the cortex of WT mice i.p injected with saline (Sal; n=6) or LPS-low (0.1mg/Kg; n=6) or APP/tTA mice i.p injected with saline (Sal; n=5) or 0.1mg/kg (LPS-low; n=6). Data analysed with 2-way ANOVA and a Tukey’s multiple comparison test. (I) RT-PCR analysis of the mRNA expression of the proinflammatory cytokines IL-1β, IL-6 and TNFα in the cortex and hippocampus of WT mice pre-treated with saline (Sal; n=5) or LPS-low (0.1mg/Kg; n=6) or of APP/tTA mice pre-treated with saline (Sal; n=4) or 0.1mg/Kg (LPS-low; n=6). IL-1β analysed with 2 way-ANOVA and Sidak’s multiple comparison test. IL-6 and TNFα analysed with multiple Mann-Whitney tests. (J) Representative images of the cortex (K) and CA1 area of WT and APP/tTA mice pre-treated with Sal or LPS-low and off DOX for 16 weeks. Scale bar 25μm. White arrows indicate Dectin-1^+^ cells (black) and plaques stained with congo red (red) (L) Dectin-1^+^ density (dectin-1^+^ cells / total area in mm^2^) in the cortex of WT mice pre-treated with saline (sal; n=6) or 0.1mg/Kg (LPS-low; n=6) or of APP/tTA mice pre-treated with saline (Sal; n=5) or 0.1mg/Kg (LPS-low; n=6). Data analysed with multiple Mann-Whitney test. (M) Dectin-1+ density (dectin-1+ cells / total area in mm2) in the CA1 area of the hippocampus of WT mice pre-treated with saline (Sal; n=6) or 0.1mg/Kg (LPS-low; n=6) or of APP/tTA mice pre-treated with saline (Sal; n=5) or 0.1mg/Kg (LPS-low; n=6). Data analysed with 2-way ANOVA test. Data shown represented as mean±SEM. Statistical differences *p<0.05, ***p<0.001.

We next studied if inflammatory cytokines might be associated with the observed increase in Aβ plaques. Pre-treatment with 3x LPS-low does not increase the mRNA expression of the proinflammatory cytokines *il-1β*, *il-6* and *tnfα* in the hippocampus or cortex of WT and APP/tTA mice **(Figure 7I)**. This is likely related to the very early stage of pathology being analysed, since the expression of these cytokines is unchanged in APP/tTA mice when compared with WT **(Figure 7I)**.

We studied the association of changes in the nascent plaque pathology with the local activation of microglial cells, focusing on dectin-1^+^ cells. Our results show that pre-conditioning with 3x LPS-low in APP/tTA mice triggers a significant increase in the density of dectin-1^+^ cells in the cortex **(Figure 7J,L)** and CA1 **(Figure 7K,M)** compared to WT mice. Altogether, these results support that inflammatory pre-conditioning accelerates the onset of Aβ pathology and the associated microglial activation.

## DISCUSSION

Systemic challenge with bacterial LPS has been widely used as an *in vivo* model of sickness behaviour (Cunningham et al., 2007, 2005; Teeling et al., 2007). Here we showed that this induction of sickness behaviour, and its associated inflammatory activation, is dose dependent with LPS-low triggering a very mild response when compared with LPS-high. Our results using LPS-high agree with the classical paradigm of systemic and central inflammation, characterised by an increased production of IL-1β, IL-6 and TNFα (Murray et al., 2011; Singh and Jiang, 2004; Verma et al., 2006). Within the hippocampus, mice dosed with LPS-high show a high concentration of TNFα and IL-1β in the brain and these cytokines have been shown to work synergically in triggering sickness behaviour and inflammation (Bluthé, Laye, et al. 2000). However, LPS-low failed to induce these changes, in turn driving increased proliferation of microglia under non-inflammatory conditions. Previous studies reported an increased proliferation of microglial cells after systemic LPS (Fukushima et al., 2015; Furube et al., 2018; Tejera et al., 2019), although the dose, timing, and localisation of this response was not fully elucidated. Previous studies had indicated that systemic LPS could trigger microglial proliferation independently of the dose, with high doses inducing increased cycling (Cazareth et al., 2014; Chen et al., 2012; Furube et al., 2018; Tejera et al., 2019). We did not observe the proliferative response at our high dose during the analysed time window, a period of time sufficient to resolve any low-grade inflammation induced by LPS-low but still premature to escape the initial inflammatory wave in LPS-high. It could be hypothesised that proliferation could occur in the LPS-high group after an extended resolution phase, in a model whereby proliferation is subsequent to inflammation (Riester et al., 2020). However, and for the purpose of our global aims, exploring these long-term effects would have difficulted any protocol of repeated systemic challenges, not allowing a clean avoidance of the post-challenge effects before a second or third dose being placed into the system.

The CSF1R-PU.1-C/EBPα axis is one of the main pathways that has been associated with microglial proliferation, and is required for microglial survival (Elmore et al., 2014; Gomez-Nicola et al., 2013; Olmos-Alonso et al., 2016). The increased local production of cytokines in response to LPS could be regulating microglial activation and proliferation (Monif et al., 2016), via the CSF1R pathway. Research using hematopoietic stem cells has shown that inflammatory cytokines, such as IL-1β and TNFα, can upregulate the expression of *Pu.1* (Etzrodt et al., 2019), a transcription factor that can control microglial proliferation by stimulating CSF1R expression (Celada et al., 1996). Our results using pharmacological inhibition of CSF1R demonstrate the mechanistic involvement of this pathway in controlling the LPS-induced microglial proliferation, although it is still unclear if the initial trigger is direct exposure to CSF1/IL34 after LPS challenge, or instead an indirect activation via PU.1.

Immune memory is a core concept of the adaptive immune system but also key for myeloid cells (Neher and Cunningham, 2019; Netea et al., 2016, 2015). LPS can trigger immune tolerance in microglial cells: a second dose of LPS in mice triggers a suppression of IL-1β release in the brain for up to 32 weeks (Schaafsma et al., 2015). Similarly, while a single dose of LPS could trigger immune priming one day later, four consecutive doses of LPS spaced one day apart triggered immune tolerance in the brain (Wendeln et al., 2018). Of note, these findings used consecutive daily doses of LPS, and this would assume that subsequent doses arrived during the inflammatory wave of previous doses (hours), instead of allowing for inflammatory resolution (days) to take place. We aimed to use the LPS-low dose to induce several rounds of microglial proliferation within a low-grade inflammatory response, but we ensured mice were fully recovered from every challenge before subsequent doses arrived. Our results show that spacing three doses of LPS-low by a week avoid a robust induction of tolerance, only showing a mild tolerance response after the third dose, reaffirming the key role of the timing of these challenges. This approach induces the expression of Dectin-1 in microglia after one dose of LPS-low, one of the markers of DAMs relevant for the pathophysiology of AD (Keren-Shaul et al., 2017). These results are in agreement with *in vitro* studies on human monocytes showing that LPS promotes the expression of dectin-1 (Rogers et al., 2013). Since we did not detect a cumulative increase in dectin-1^+^ cells after weekly challenge our findings present two potential interpretations. First, the dectin-1^+^ cells identified at week 1 may be the same cells observed at week 3, exhibiting a decline over time, suggesting that only the initial dose of LPS-low had an effect. Alternatively, another interpretation posits that each LPS-low dose induces a transient increase in dectin-1^+^ cells, which resolves before the subsequent challenge, potentially to be reactivated following successive doses. In both cases, we would need more specific phenotyping of these microglia in order to understand whether the induction of dectin-1^+^ is transient or permanent after LPS challenge, or whether it is connected to proliferation.

The links between systemic inflammation and brain dysfunction are wide and far reaching. Elderly individuals with high levels of circulating IL-6 often show a reduced hippocampal volume and increased cognitive decline (Marsland et al., 2008). Circulating cytokines have been linked with neurodegenerative diseases such as PD and AD (Holmes et al., 2009; Rathnayake et al., 2019), indicating that systemic inflammation can modify the trajectory of neurodegenerative diseases. In addition, during neurodegeneration, LPS causes an exacerbated inflammatory response in the brain, followed by increased cognitive decline and acceleration of the disease (Skelly et al., 2013). Mouse models of AD-like pathology show higher production of IL-1β when administered with LPS, supporting the hypothesis that ongoing neurodegeneration primes microglia to further inflammatory challenge (Lee et al., 2008; Tejera et al., 2019). However, few studies have studied this relationship in the inverted order: can the exposure to systemic inflammation during an otherwise healthy early life prime microglia and affect the onset of AD? Here we took advantage of the repressible nature of the APP/tTA model to drive systemic low-grade inflammation during healthy adult life, evidencing that the pre-conditioning of microglial cells with 3x LPS-low modified the onset of AD-like pathology, triggering a higher accumulation of Aβ plaques.

Previous studies, using different models of neurodegeneration, have shown that using a secondary stimulus of LPS or live bacterial challenges, negatively impacts the disease (Chouhan et al., 2022; Cunningham et al., 2009; Sly et al., 2001). Our results now support that the inverse relationship is also possible, perhaps a scenario reflecting the trajectory seen in humans more optimally: humans are very regularly exposed to low-grade inflammation during adult life, preceding the onset of AD. Our data suggests that the onset of Aβ pathology is not directly influenced by an exacerbated inflammatory response, despite the increase in dectin-1^+^ seen after 3x LPS-low challenge, or by influencing the expression of APP. Instead, we speculate this might be triggered by a change in the processing of APP (Lee et al., 2008), associated by the increase in dectin-1^+^ cells. A recent study from our lab showed that a subpopulation of dectin-1^+^ showed senescence markers and were associated with an increase in amyloid pathology (Hu et al., 2021). In our model, LPS-low might be triggering this subpopulation of dectin-1^+^, possibly by an increase in proliferation, shown to negatively impact the onset of the disease (Hu et al., 2021; Mancuso et al., 2019; Olmos-Alonso et al., 2016). However, future studies should address this by studying the dectin-1^+^ population and other DAM markers under these circumstances, and how they directly affect amyloid pathology.

Taken together we demonstrated that systemic LPS has a dose and time-dependent effect on brain inflammation and on the proliferation of microglia. LPS-high (1mg/kg) triggers an acute peripheral and central inflammatory response that lasts at least three days and is accompanied by sickness behaviour directly or through mediators that activate microglial cells. On the contrary, LPS-low (0.1mg/kg) promotes a CSF1R-dependent microglial proliferation without inducing an inflammatory response in the brain. Repeated systemic challenges with low-grade inflammation, timed to avoid a robust tolerance response, have long-lasting consequences on the onset of amyloid pathology, accompanied by the induction of dectin-1^+^. These findings have significant relevance for the understanding of AD in humans and how low-grade inflammation experienced during early healthy life determines the onset of the pathology.

## Acknowledgements

We thank the Southampton Biomedical Research Facility for assistance with animal breeding and maintenance. We thank Georgina Dawes for technical assistance. We thank Dr. Joanna Jankowsly for the line 102 mice.

## Data availability

The data that support the findings of this study are available from the corresponding author upon reasonable request.

## Ethics approval

All experimental procedures were approved by a local ethical review committee and conducted in accordance with relevant personal and project licenses under the UK Animals (Scientific Procedures) Act (1986).

